# Spatio-temporal patterns of whiting (*Merlangius merlangus*) in the Adriatic Sea under environmental forcing

**DOI:** 10.1101/2023.08.01.551447

**Authors:** Emanuele Asciutto, Federico Maioli, Chiara Manfredi, Alessandra Anibaldi, Jacopo Cimini, Igor Isailović, Bojan Marčeta, Michele Casini

## Abstract

Understanding how environmental factors affect species distribution is crucial for the conservation and management of marine organisms, especially in the face of global changes. Whiting (*Merlangius merlangus*) is a demersal temperate fish, considered a ‘relict species’ in the Adriatic Sea. Despite its significance to commercial fisheries in the region, the specific drivers behind its spatial and temporal patterns have not been thoroughly examined. Here, we fitted a set of Generalized Linear Mixed Effects Models to data collected in the Northern and Central Adriatic from 1999 to 2019 during the Mediterranean International Trawl Survey to investigate the potential influence of depth, seafloor temperature and bottom dissolved oxygen on the annual biomass density and spatial distribution of whiting. Our results showed that depth, and to a lesser degree temperature and oxygen, are important predictors of whiting distribution in spring-summer, with preferences for depths of ∼ 45 m, temperature of ∼ 15.4 °C and dissolved oxygen > 5.5 ml L^-1^. We predicted a persistent core area of distribution in front of the Po River Delta, in the Northern Adriatic Sea, while the density progressively declined towards the Central and Southern Adriatic Sea along the Italian coast. Additionally, the temporal trend exhibited high fluctuations over the years, occurring in cycles of 3 to 4 years. Finally, by comparing the biomass density estimates obtained under optimum conditions with those derived from the actual values for each variable, our analysis revealed that temperature had a pronounced and general impact on biomass density in the northern survey area (prediction revealed a ratio of density reduction of approximatively a half), while oxygen displayed a minor and more localized influence. This work deepens the current knowledge about the ecology of whiting in the Adriatic Sea and provides support for the conservation and management of this species.

## 1 Introduction

Tracking and understanding the effects of environmental factors on the distribution of marine organisms is crucial for the management of species of conservation or commercial interest [1,2], especially in light of major global changes. As a matter of fact, marine species are quickly reorganizing due to climate change [3], even faster than species on land [4]. Generally, marine species are expected to shift poleward [5,6], but also towards deeper and colder areas where local geographical barriers impede latitudinal shifts [7,8]. However, departures from these expectations might occur because of individual species responses, local and regional factors [9], differences in species interactions and stochastic factors [10,11]. Moreover, organisms are likely exposed to diverse aspects of climate change, e.g. changes in sea water level and temperature, acidification, deoxygenation, modifications in rivers run-off, and change in frequency and strength of adverse weather conditions [12]. This picture is further complicated by the potential effects of other anthropogenic drivers including eutrophication [13], fishing pressure [14,15], pollution [16], and habitat degradation [10], which may interact with climate change at multiple scales.

The impact of global changes on marine species is particularly severe in areas where range shifts are physically constrained, such as in semi enclosed areas [17]. The Adriatic Sea is a semi-closed basin of the Mediterranean Sea which has experienced significant environmental fluctuations and ecosystem degradation linked to the combined effects of climate change, increasing fishing effort and eutrophication in recent decades [18–21]. The marine species living in this basin seem to be particularly vulnerable to environmental disturbances (e.g., long term temperature change, modification of hydrological regime, changes in production and food availability; [22,23]).

Here, we employed data from two decades of fishery-independent surveys to investigate the effects of environmental drivers on whiting, *Merlangius merlangus,* in the Northern-Central Adriatic Sea where this species constitutes a common species of the demersal community [24]. Whiting is a cold-temperate demersal fish widely distributed in the North-East Atlantic Ocean [25], whereas in the Mediterranean Sea [26] and in the Black Sea [27] it represents a ‘relict species’ which likely migrated from northern waters during the last glacial period [24,28]. In the Mediterranean Sea whiting is particularly abundant in the Northern-Central Adriatic Sea [29], where it is commonly found near muddy or sandy bottoms down to depths of 100 m [30–32]. Some specimens can also be found along the central and southern Italian coast [33]. This species has a relevant role in the fisheries of several countries, and it is included into the list of species of commercial and conservationist interest in the Mediterranean and Black Sea by the General Fisheries Commission for the Mediterranean (GFCM). According to data collected up to 2017, this species accounted for 3.2% of the total bottom trawl fishery landings in the Adriatic Sea (on a total of 28,800 tons) [34].

Despite the widespread distribution of whiting and its commercial relevance in the Northern-Central Adriatic Sea, many aspects of the ecology of this species have not been analytically investigated so far in this basin. In the present study we use spatio-temporal species distribution models with the aim of a) characterizing the spatial and temporal variation in whiting biomass and b) investigating the effect of the major environmental factors on its spatio-temporal dynamics.

## 2 Materials and methods

### 2.1 Study area

The study area is represented by the Northern-Central Adriatic Sea, a semi-enclosed basin corresponding to the FAO - Geographical Sub-Area (GSA) 17 [35] located in the Central Mediterranean, between the Italian peninsula and the Balkans. The basin is a shallow and mainly eutrophic sea with clear morphological differences along both the longitudinal and the latitudinal axes. Most of the basin is characterized by a wide continental shelf where depth gradually increases from North to South. Sea bottom is mostly covered with recent muddy and coastal sandy sediments supplied by rivers and dispersed by marine current, while the offshore seabed in the Northern Adriatic is characterized by relict sand [36]. The eastern coast is rocky and articulated with many islands whereas the western one is flat, alluvial, and characterized by heavy river runoff. The Po and the other northern Italian rivers contribute to approximately 20% of the whole Mediterranean river runoff [37] and introduce large fluxes of nutrients [38], making this basin the most productive area of the Mediterranean Sea [39] and consequently one of the most intensively fished areas in Europe [40]. The general pattern of water circulation is cyclonic with water masses inflow from the Eastern Mediterranean along the eastern side of the Adriatic Sea and outflow along the western side. Being a continental basin, the Adriatic Sea circulation and water masses are strongly influenced by river runoff and atmospheric conditions that affect salinity and temperature [41]. Pronounced seasonal fluctuations in environmental forcing, cause high seasonal variation in coastal waters, where bottom temperatures, range from 7 °C in winter to 27 °C in summer [42]; in the deeper areas the thermal variability is reduced, with values ranging between 10 °C in winter and 18 °C in summer at a depth of 50 m [42]. A well-developed thermocline is present in spring and summer [41] which, under suitable marine weather conditions (i.e., high temperatures, extended period of calm sea, freshwater inflow, stratification) can cause exceptional algal blooms and extended hypoxia and anoxia phenomena that damage the demersal and the benthic communities [43].

### 2.2 Data source

#### 2.2.1 Survey data

Biomass density data of whiting (i.e., estimated as the ratio between the total weight of fish caught and the area swept in each haul, expressed as kg km^-2^) were obtained from standardized experimental trawl hauls carried out in the GSA 17, within the frames of the Mediterranean International Trawl Survey (MEDITS). This survey started in 1994 at the European Mediterranean level, with the aim to collect data on demersal communities to describe their distribution and demographic structure [44,45]. The MEDITS covers the whole GSA 17 since 1996 and it is jointly performed by the Laboratory of Marine Biology and Fisheries of Fano (Italy), the Institute of Oceanography and Fisheries of Split (Croatia) and the Fishery Research Institute of Slovenia. Between 88 and 247 trawl hauls were performed annually, typically during the late spring-summer season (May-September) using an experimental bottom trawl with 20 mm of stretched mesh cod-end. Hauls were located from 10 to 500 m of depth, following a random-stratified sampling scheme where strata were defined according to depth. Further details on sampling procedures, data collection and analysis can be found in the MEDITS handbook [46]. In the present study, we included trawl hauls performed between 1999 and 2019 because of the lack of seafloor dissolved oxygen data for earlier years. In order to have a more seasonal-homogeneous dataset, we included only the hauls performed in May-September and removed those conducted in the autumn-winter period (i.e., October-December) (S1 Fig).

#### 2.2.2 Environmental variables

In order to investigate major changes in the environmental covariates and potentially concurrent changes in whiting biomass density, we specifically focused on three key factors: depth, seafloor temperature (hereafter temperature; T), and seafloor dissolved oxygen (hereafter dissolved oxygen; DO). These factors were selected based on previous knowledge [47] and data availability. T and DO data used in this study were obtained from the E.U. Copernicus Marine Service Information [48,49] extracting the monthly mean of each variable. Hence, to integrate the environmental data with the survey data, we associated each survey haul with the nearest spatial point where the variables were available, as well as the nearest corresponding time reference for the haul.

### 2.3 Statistical analysis

#### 2.3.1 Quantifying local environmental trends

To assess and visualize potential changes in the environmental drivers, we estimated the mean values across the study area and years as well as their local rate of change. To achieve this, we constructed a 2 km^2^ grid (UTM 33N projection) encompassing the entire survey area, and matched the environmental covariates as described is section 2.2.2. Additionally, depth data was retrieved from the EMODnet Bathymetry project, https://www.emodnet.eu/en/bathymetry, funded by the European Commission Directorate General for Maritime Affairs and Fisheries. Thereafter, for temperature and dissolved oxygen, we employed a linear model for each grid cell, adopting the following form:

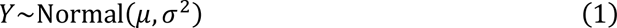

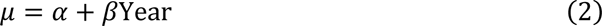

Here, *Y* denotes the environmental covariate value at the cell, μ denotes the mean of the cell, and *σ* represents the standard deviation. The intercept is denoted by *α*, and the coefficient related to year is represented by *β*. The local rate of change can be derived from the *β* slope. To facilitate interpretation, we presented the local (i.e., the grid cell) rate of change as decadal trends.

#### 2.3.2 Spatio-temporal models

The spatio-temporal distribution of whiting was analysed using geostatistical Generalized Linear Mixed Effects Models (GLMMs) fitted by maximum marginal likelihood using the sdmTMB R package (version 0.0.21.9;[50]) in R (version 4.0.3; [51]). Biomass density was modelled with a Tweedie distribution and a log link function to accommodate both zeros and positive continuous values [52]:

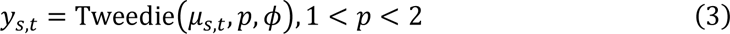

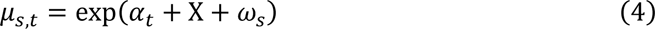

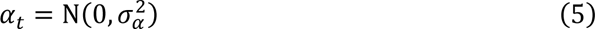

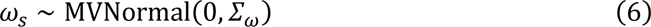

Where *y****_s_****_,t_* is the biomass density (kg km^-2^) at point ***s*** in space and *t* in time, 𝜇_𝑠,𝑡_the mean biomass density, *p* the Tweedie power parameter and 𝜙 the Tweedie dispersion parameter. The parameter 𝛼_𝑡_ is the random intercept by year *t* with variance 𝜎_𝑎_. The parameter *X* represents the design matrix of the covariates (which are specified in Table 1). The 𝜔_𝑠_ parameter represents a spatial random field. Spatial random fields are assumed to be drawn from a multivariate Gaussian distribution with covariance matrix 𝛴_𝜔_ and constrained by a Matérn covariance function. Following Essington et al., 2022 [53], we intentionally omitted the spatiotemporal random fields (𝜀_𝑡_) to specifically focus on examining the relationship between inter-annual changes in biomass density and variations in T and DO. The inclusion of spatiotemporal random fields introduces an additional complexity when attempting to disentangle the individual effects of T and DO on biomass density changes [53].

**Table 1.** Models’ formulations and model selection results. Df are the degrees of freedom. DoY is the day of the year, T is the temperature and DO is the dissolved oxygen. AIC represent the scores of the Akaike Information Criterion.

| Model | Formula | df | AIC | ΔAIC |
| --- | --- | --- | --- | --- |
| Model 0 | Depth + Depth<sup>2</sup> + DoY + DoY<sup>2</sup> | 10 | 13603.43 | 196.56 |
| Model 1 | Depth + Depth<sup>2</sup> + DoY + DoY<sup>2</sup> + T | 11 | 13516.90 | 110.03 |
| Model 2 | Depth + Depth<sup>2</sup> + DoY + DoY<sup>2</sup> + T<br>+ T<sup>2</sup> | 12 | 13418.62 | 11.75 |
| Model 3 | Depth + Depth<sup>2</sup> + DoY + DoY<sup>2</sup> +<br>DO | 12 | 13518.85 | 111.98 |
| Model 4 | Depth + Depth<sup>2</sup> + DoY + DoY<sup>2</sup> + T<br>+ DO | 13 | 13467.52 | 60.65 |
| Model 5 | Depth + Depth<sup>2</sup> + DoY + DoY<sup>2</sup> + T<br>+ T<sup>2</sup> + DO | 14 | 13406.87 | 0.00 |

#### 2.3.3 Model specification and model selection

All models in our analysis accounted for fixed effects of day of year (DoY) and depth, modelled as linear plus quadratic terms. Models 1 to 5 further incorporated different combinations or specifications of T and DO as covariates (see Table 1). DO was included as a breakpoint (BP) effect, which assumes that there is no effect of DO on biomass density above a certain threshold, while below this threshold, the effect is linear [53]. All covariates were centered and scaled by their standard deviations to aid convergence and facilitate comparison of coefficients [54]. We used the Akaike Information Criterion (AIC, [55]) to compare the performance of the five models with different combinations of covariates.

#### 2.3.4 Spatio-temporal predictions of biomass density

To evaluate changes in whiting biomass density over space and time, we utilized the best-fitting model to predict onto the 2 km^2^ grid. We measured the standard errors associated to each grid cell prediction by sampling 500 simulations from the joint precision matrix. By multiplying the area with the biomass density and summing up the results across the entire study region, we derived an overall model-based temporal index of the biomass density for the Northern-Central Adriatic Sea. This index represents the predicted biomass density per year and was further normalized using the first year of the data used (i.e., 1999).

## 3 Results

### 3.1 Summary of environmental variables

The northern part of the Adriatic basin is characterized by shallow waters (mean depth of ∼ 40 m) with a weak bathymetric gradient along the longitudinal axis (Fig 1a-b). The central area is characterized by meso-Adriatic depressions (Pomo Pits), where the depth decreases to ∼ 270 m. In the Northern-Central Adriatic, high seafloor summer temperatures (T) were registered in the Italian shallow coastal areas (∼ 25°C), while colder waters were in deeper zones and along the Croatian coast (∼ 15°C) (Fig 1c). Over the study period, T increased by an average of ∼ 0.30 °C per decade across the entire basin, but with more pronounced warming offshore the Po River Delta, south of the Pomo Pits and along the central and southern Italian coast (up to ∼ 1 °C per decade). In contrast, the Gulf of Venice and the central Croatian coast showed a decrease in T of as high as ∼ 0.5 °C per decade (Fig 1d). The bottom dissolved oxygen (DO) was lower along the central-southern Adriatic (∼ 5.4 ml/L^-1^) (Fig 1e). The highest values were registered north and south of the Po River Delta and in the Croatian channels (∼ 6.0 ml/L^-1^). DO displayed, along the study period and across the study area, a decrease by an average of 0.02 ml/L^-1^ per decade, with a general increase in the Northern Adriatic (i.e., areas below Po River Delta, ∼ 0.15 ml/L^-1^) and a slight decrease in the south-western areas (∼ −0.05 ml/L^−1^). A decline in DO was also observed in small areas just below the Istrian Peninsula and along the shore of the Gulf of Venice.

**Fig 1.**
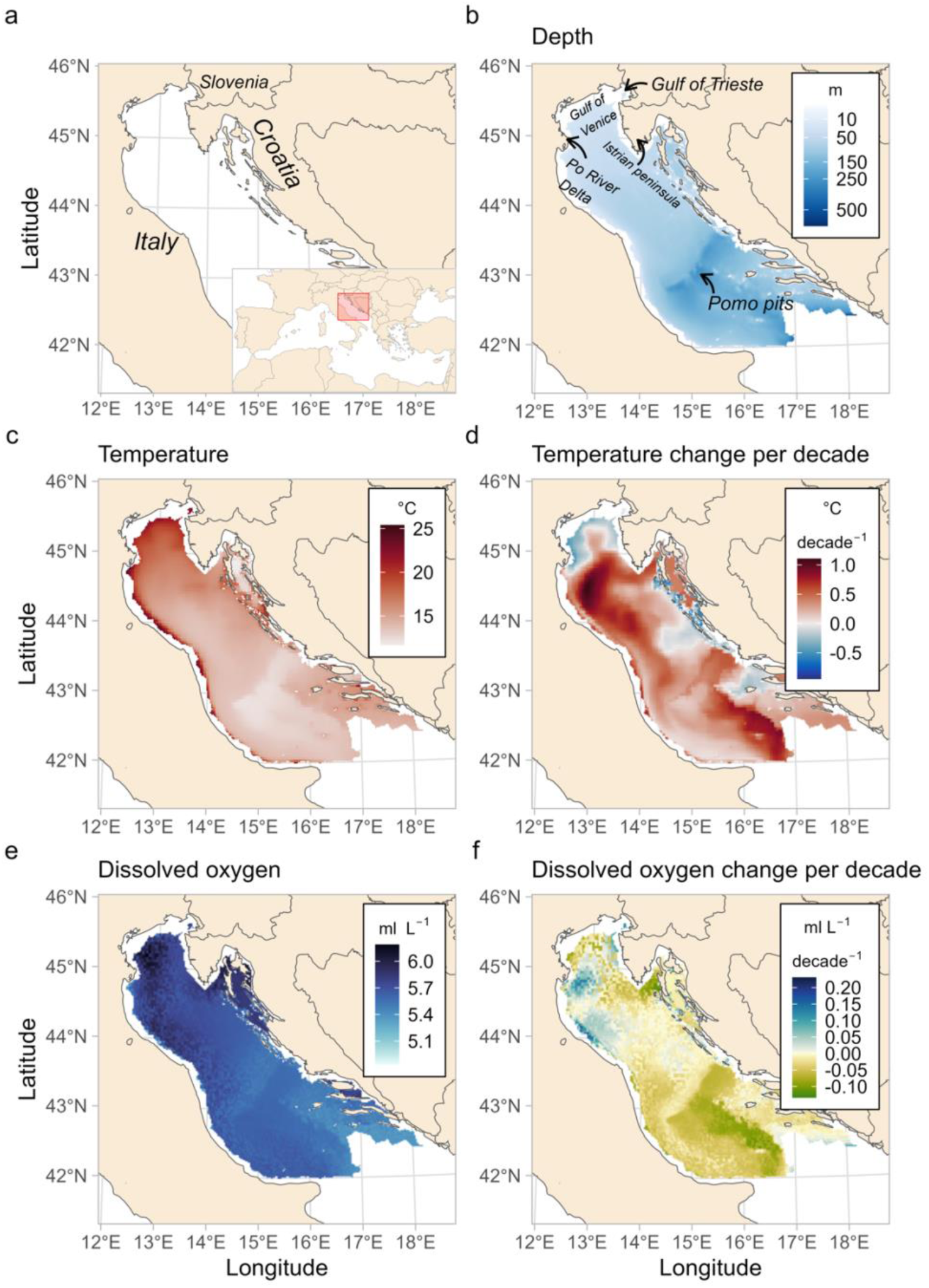
Study area (a), depth (b), mean temperature (c) and mean dissolved oxygen (d) conditions (June-August) between 1999-2019, and predicted decadal local trends for temperature (d) and oxygen (f). The local rate of change was calculated for each grid cell (2 km^2^).

### 3.2 Best fitting model and effects of environmental variables

Among the models we tested (listed in Table 1), Model 5 achieved the lowest AIC score, indicating it as the best-fitting model. Model 5 included T, T^2^ and DO, in combination with DoY, DoY^2^, depth, depth^2^, spatial random effects (𝜔_𝑠_), as well as a random year intercept (𝛼_𝑡_). The estimated coefficients for Model 5 showed a strong effect of depth, and to a lesser extent of DO and T, even after accounting for spatial random effects (S2 Fig). Despite the presence of some spatial clustering, the residual diagnostics of our model looked satisfactory, indicating that the model adequately captured the relationship between the environmental factors and the distribution of whiting populations. More information about model diagnostics can be found in the Supporting Information (S3 and S4 Figs). We calculated optimal values for each environmental variable and examined the biomass density variation associated with these values. Taking into account the minimum and maximum recorded values for T (9 °C and 28 °C, respectively), the minimum recorded value for DO (4.7 ml L^−1^), and the shallowest depth considered in the MEDITS protocol (10 m), we evaluated the lower and higher ranges for each variable (i.e., the variation in biomass density from the minimum and maximum value to the optimum). Based on our analysis, an optimal depth of approximately 45 m (Fig 2a) was estimated for whiting. The mean biomass density exhibited an increase of 1.8 times from shallow depths of 10 m towards the estimated optimum. However, it dropped close to zero at depths exceeding 110 m (Fig 2a). The predicted mean biomass density peaked at an intermediate T of 15.4 °C (Fig 2b). The biomass density displayed an increase of 2.4 times from the cold T of 9 °C to the optimum, while it showed a decrease of −1.9 times from the optimum to highest T of 28 °C. Additionally, we estimated a DO threshold of approximately 5.5 ml L^−1^ (Fig 2c) with a 95% confidence interval of [5.41, 5.62]. Under suboptimal conditions of 4.7 ml L^−1^, there was a variation in biomass density of −3.1 times.

**Fig 2.**
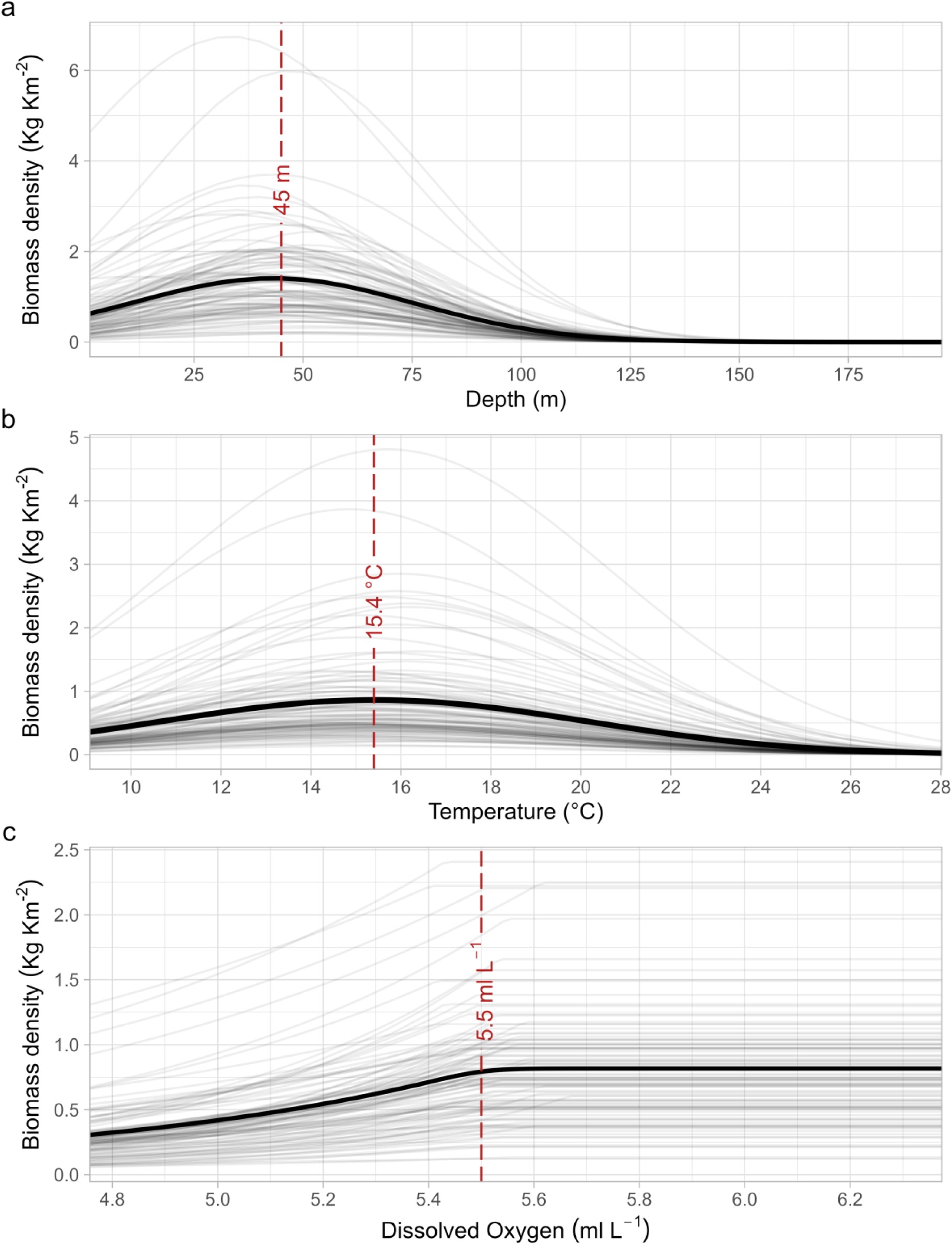
Marginal effects of depth (a), temperature, T (b) and dissolved oxygen, DO (c). Gray lines represent random draws from the joint precision matrix (with n of simulations = 100) while black lines represent mean values. Depth and T were modelled as a quadratic function to allow for intermediate optima, while DO was modelled as breakpoint function. The red dashed line marks the optima of depth and T, and the breakpoint of DO.

### 3.3 Spatial and temporal variations in biomass density

Whiting occupied nearly half of the Northern-Central Adriatic Sea (Fig 3a and S5 Fig). Predicted high densities (up to ∼ 600 kg km^2^) indicated that most of the whiting inhabits the northern part of the basin, with a core area located between the Po River Delta and the Istrian peninsula. Predicted intermediate values of biomass densities (∼ 100-150 kg km^−2^) resulted to be spread around the core area, in the central part of the basin, and along the Italian coast, while the offshore area in the central part of the basin displayed lower densities (as low as ∼ 10 kg km^−2^). Predicted densities reached nearly 0 in the offshore central and southern part of the basin where the depths exceed 140 m, and along the central and southern Croatian coast. During the entire time series whiting occupied broadly the same areas. The highest standard error values were estimated in the southern part of the basin (Fig 3b). The spatial biomass density predictions for each year of the time-series are available in the Supporting Information (S5 Fig). The overall temporal biomass density index, based on the best-fitting model, exhibited significant fluctuations during the period under investigation (Fig 3c). These fluctuations manifested as consecutive peaks and valleys occurring approximately every 3-4 years.

**Fig 3.**
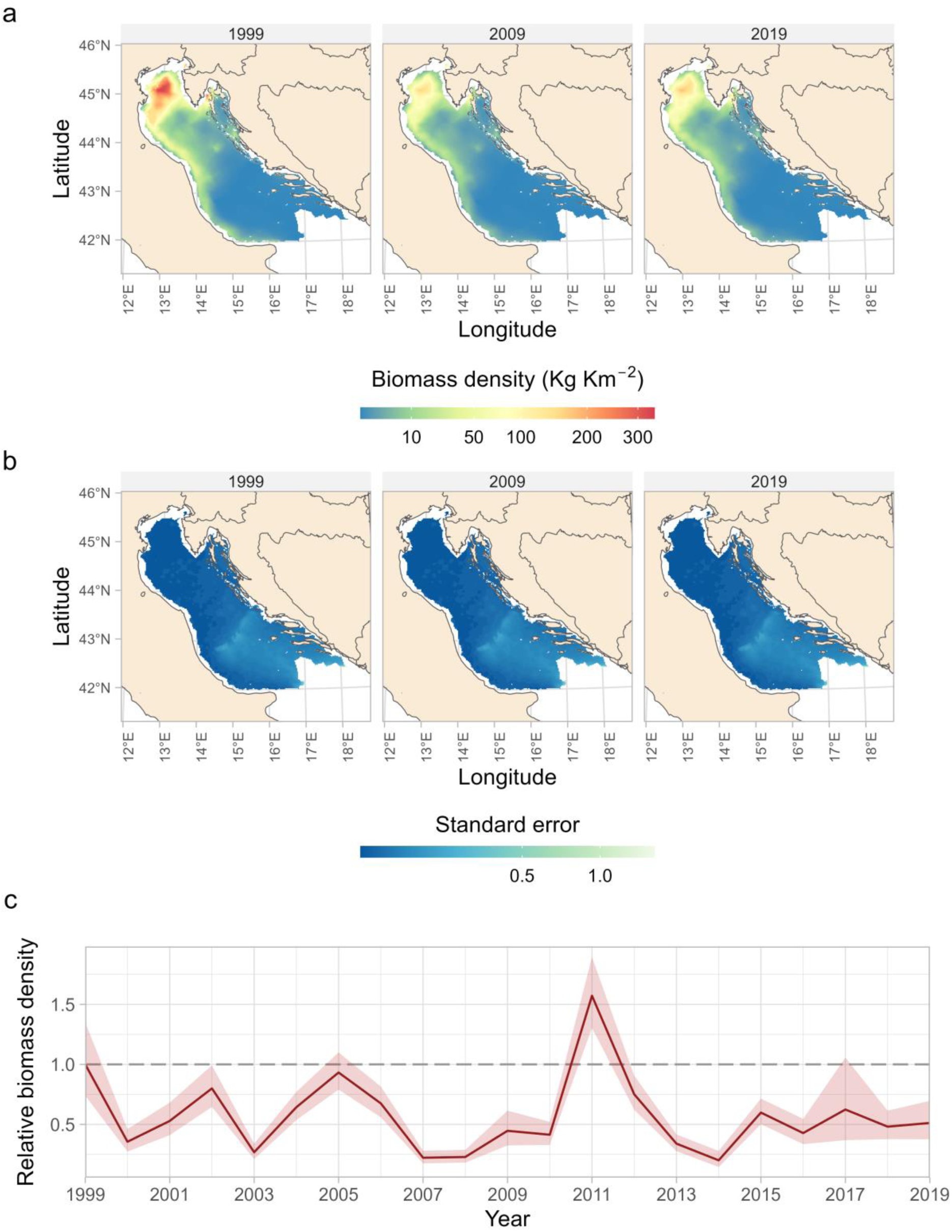
Predicted spatio-temporal changes of whiting biomass density (Kg Km^−2^) (a) with associated standard error of the mean (b). Relative temporal biomass density index normalized to the first year (c). All predictions were made over a 2 km^2^ prediction grid. For the predictions of all the years see S5 Fig in Supporting Information. In panel (c) the values have been standardized to the initial year of the survey, namely 1999, which is represented by the dashed line. The shades around the mean are 95% confidence intervals.

### 3.4 Predicted biomass density under different scenarios

To further assess the influence of T and DO on whiting, we conducted separate analyses comparing predicted biomass density under different environmental scenarios. For T, we compared two scenarios: one assuming the estimated optimum T at all sites, and the other incorporating the actual T values (Fig 4). Our analysis revealed significant effects of T, particularly in the northern part of the study area (Fig 4). Here, biomass densities were predicted to be reduced by half compared to the expected levels under optimum T conditions.

**Fig 4.**
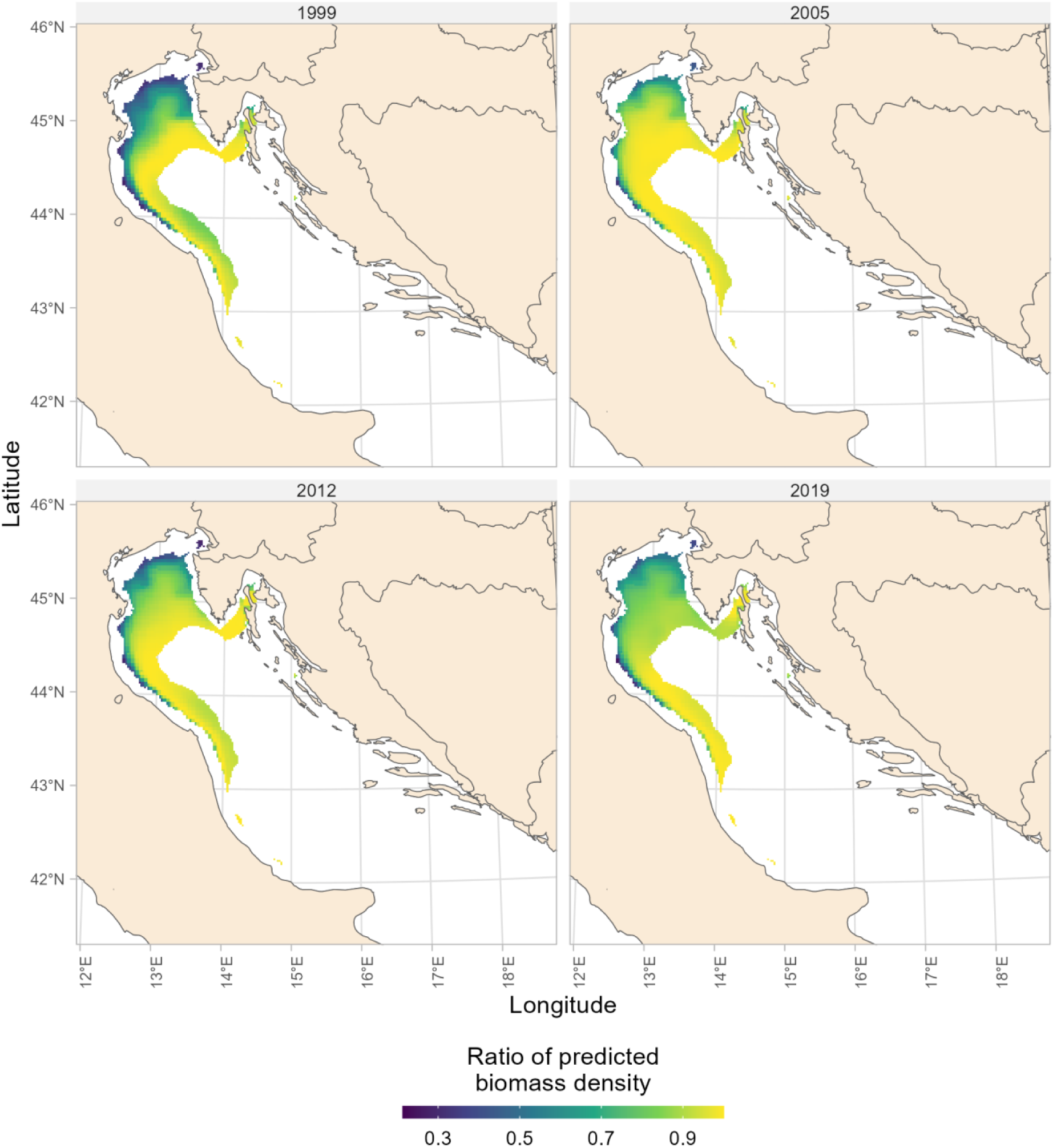
Map of the the proportional reduction in whiting biomass density attributed to variations in temperature (T). The proportional reduction in whiting biomass density is computed as the ratio of predicted densities at the estimated optimal T compared to the actual T, based on the best fitting model. Only cells that cumulatively reached 90% of the whiting biomass density are displayed. The years are displayed considering an interval of 7 starting from the first. All predictions were made over a 2 km^2^ prediction grid.

Similarly, we explored the impact of DO levels on whiting by comparing predicted biomass density under two distinct scenarios. The first scenario assumed that DO at all sites exceeded the estimated threshold, while the second scenario utilized the actual extracted values of DO (Fig 5). Our analysis highlighted localized effects of DO, particularly in the coastal areas south of the Po River Delta, extending to the northern study area and along the western side of the Istrian Peninsula. In these areas, biomass density was predicted to decrease to approximately one-quarter to half of the expected levels observed at the estimated threshold of DO conditions, depending on the specific location.

**Fig 5.**
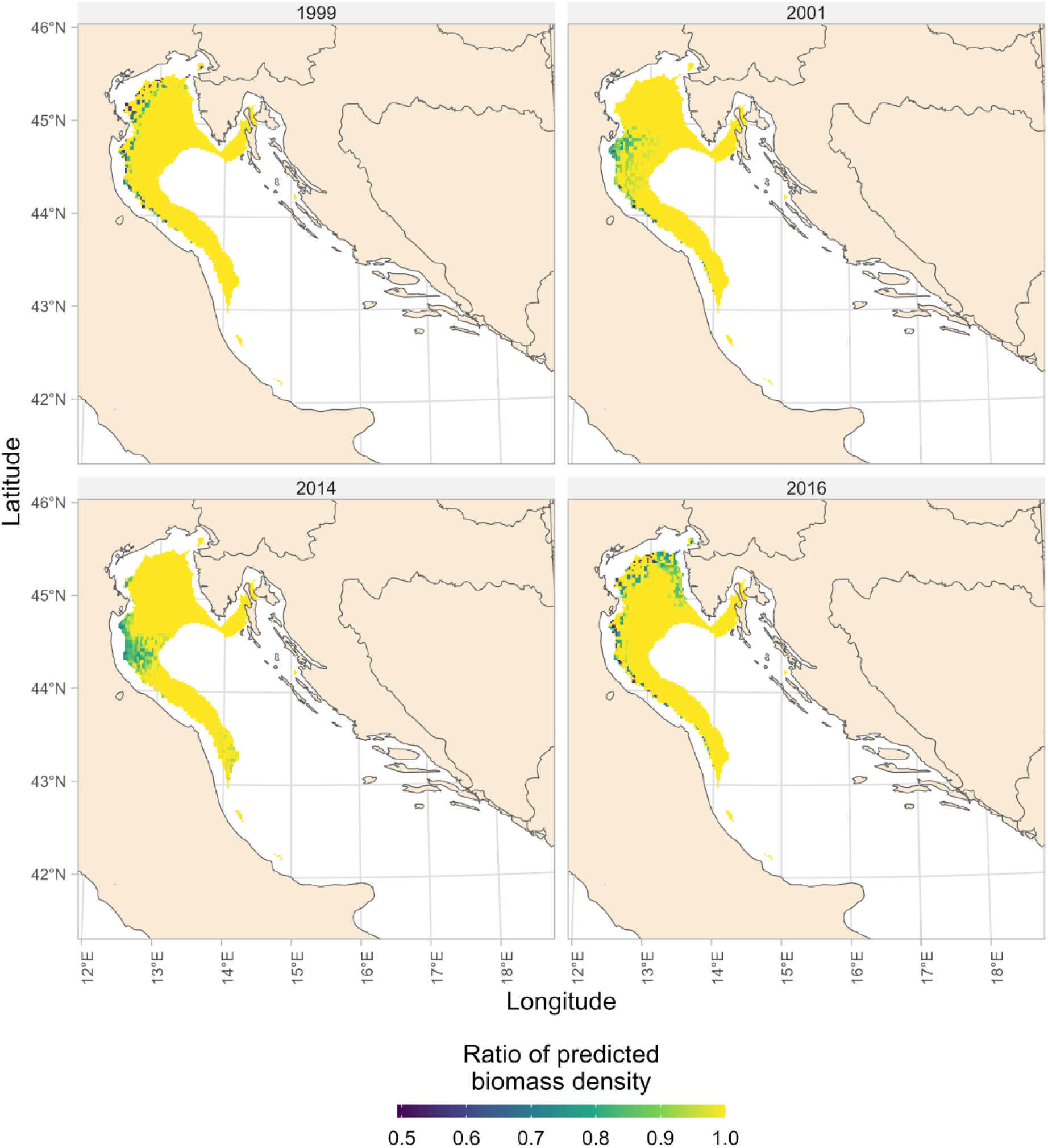
Map of the the proportional reduction in whiting biomass density attributed to variations in dissolved oxygen (DO). The proportional reduction in whiting biomass density is computed as the ratio of predicted densities at the estimated DO threshold compared to the actual DO, based on the best fitting model. Only cells that cumulatively reached 90% of the whiting biomass are displayed. Only the most representative years were displayed. All predictions were made over a 2 km2 prediction grid.

## 4 Discussion

Our study provides an important step ahead towards a deeper ecological understanding of the response of a demersal species to major environmental variations. Here, by fitting distribution models to two-decades of fishery-independent survey data we show that depth, and to a lesser extent temperature and dissolved oxygen play a major role in shaping whiting spatio-temporal variations in biomass density over the Central-Northern Adriatic Sea. Here, we also provide a temporal index of the biomass density for the entire study area. This allows to visualize the predicted biomass density trend over the years. Furthermore, we compared the biomass densities predicted at the temperature and dissolved oxygen levels that are optimal for whiting with the densities observed at the actual values of each variable. By quantifying these disparities, we are able to highlight the magnitude of the impact that deviations from the optimal environmental conditions can have on the distribution and abundance of whiting in the Central-Northern Adriatic Sea.

### 4.1 Effect of environmental factors

Very little previous information is available on the effect of environmental factors on whiting distribution [56,57]. With regard to the Mediterranean Sea, only Papaconstantinou (1982) [26] analysed the distribution of this species in the Aegean Sea as a function of major environmental factors. In our study, we found whiting to preferably inhabit depths of about 45 m, becoming virtually absent at depths > 110 m. This is in agreement with previous studies in the Adriatic Sea [30,58] and consistent with other findings in other areas, such as in the North Sea [59], Black Sea [60] and the Aegean Sea [26,61].

Papaconstantinou (1982) [26] indicated the temperature range of whiting in the Aegean Sea as between 9.4 °C and 17.5 °C with preferences for temperatures higher than 10 °C. This is close to our estimates revealing an optimal temperature of ∼ 14.4 °C. Moreover, our results are in accordance with Svetovidov (1948) [62] who suggested that this species in the Black Sea migrates during winter from the colder areas to the warmer areas to find the optimal temperature range of 5 °C - 16 °C. However, the preferred temperatures of 6 - 10 °C reported for adult individuals in the Black Sea [63] as well as of 7 - 11 °C [64] found in the Fishbase database [65] are not completely in accordance with our results. The reason could be that the dataset included in our study only referred to the spring-summer period when water temperatures are higher. Further, we estimated the threshold of dissolved oxygen concentration at ∼ 5.5 ml L^−1^, below which the dissolved oxygen concentration seems to be suboptimal for whiting. The positive relationship between biomass density and oxygen might suggests a tendency for withing to avoid areas with suboptimal dissolved oxygen. Although a threshold of 1.4 ml L^−1^ (∼2 mg L^−1^) is used in most conventional applications as the level below which hypoxic conditions are met and fish stocks may start to collapse [66], as reported by two meta-analyses [67,68] more realistic sub-lethal threshold can vary by one order-of-magnitude among marine organisms. For example, cod *Gadus morhua*, a close relative of whiting, has an estimated sub-lethal threshold of ∼ 7.14 ml L^−1^ [67], even higher than our estimate. However, we recall that our threshold is estimated from in-situ observations and is therefore expected to differ from laboratory-conducted experiments. Eventually, our analysis could not exclude size-specific responses to environmental variables that could potentially explain discrepancies with literature in other areas.

### 4.2 Spatio-temporal dynamics

Our study show that whiting is mainly distributed in the northern area of the Adriatic Sea, with the core area located in the shallow waters in front of the Po River Delta. Biomass density declines towards the Central Adriatic Sea, where depth increases. The scarce, yet constant, presence of whiting along the southern Italian coast suggests that the cyclonic pattern of water circulation might carry eggs and pelagic larval stages from the northern spawning locations (also corresponding to the adults distribution area; [33]) towards these southern areas [33,69]. In the northern part of the basin, we observed an offshore area, located between the Istrian peninsula and the Italian coasts, with systematically lower densities compared to the nearby areas. This area, called “area degli sporchi” (roughly meaning “dirty area”) by fishermen, is characterized by relict sand rich in epifaunal organisms, whereas whiting seems to prefer sand-muddy sediments with scarce biocenosis [31,70]. Indeed, we predicted higher whiting densities within the continental shelf of North and Central Adriatic, mostly covered by fine sediments (muddy sand and sandy mud, [71]). Furthermore, these areas are also characterized by a scarcer density of hake (*Merluccius merluccius*), which may indicate a possible competition for the resources [24]. Moreover, within the Bristol Channel and the Severn Estuary (UK), growth and local abundance of whiting were positively linked to the presence of two preys, the brown shrimp (*Crangon crangon*) and the sprat (*Sprattus sprattus*) [72], which are also present in the northern and central areas of the Adriatic Sea [73,74]. The persistence of whiting in the same areas across years found in our study may depend also on its low tendency to make long distance movements, although the capability of some adult specimens to do so [69]. Overall, biomass density time-series was characterized by high fluctuations with peaks occurring with a frequency of 3 to 4 years, likely due to the short lifespan of this species and driven by strong recruitment events [60,75]. By comparing the biomass density estimates obtained under optimum environmental conditions with those derived from the actual values, we discovered distinct influences of temperature and dissolved oxygen on different areas within the study region. Firstly, the comparison revealed that the northern extent of the survey was primarily influenced by temperature changes. These temperature variations resulted in lower biomass density estimates, indicating that temperature could potentially act as a limiting factor in this area. The observed decrease in biomass density suggests that whiting populations are less abundant in regions experiencing higher temperatures. This finding highlights the sensitivity of whiting to temperature and suggests that temperature plays a critical role in shaping the distribution patterns of this species. In contrast, more localized areas, particularly those located south of the Po River Delta, extending to the northern part of the survey, and along the western side of the Istrian Peninsula, were affected by sub-optimal oxygen levels. These areas exhibited decreased biomass densities, indicating that whiting populations in these regions are influenced by sub-optimal dissolved oxygen levels. However, the impact of sub-optimal oxygen levels was observed within a more limited geographic extent compared to the broader temperature-related effects observed in the northern region. Overall, our study underscores the complex interplay between depth, temperature, dissolved oxygen, on the spatio-temporal dynamics of whiting, providing crucial insights into the ecological factors shaping the distribution and abundance of this demersal species in the Central-Northern Adriatic Sea.

## 5 Conclusions

Understanding how environmental factors drive the patterns of species distribution is a key ecological question and a requirement for effective management when fish species are of commercial or conservation interest. Here, we employed species distribution models to provide detailed information on the spatial ecology of whiting in the Central and Northern Adriatic Sea, one of the most commercially trawled areas of the Mediterranean. We showed that depth, and to a minor degree temperature and bottom dissolved oxygen, are important environmental factors in shaping whiting distribution in this basin. We then compared the differences in predicted biomass density between two different scenarios (i.e., optimal conditions and actual values) for temperature and dissolved oxygen. For temperature we found a decline of approximatively a half between the two scenarios, while oxygen displayed a minor and more localized influence. Taking into account the current effects of worldwide climate change, these results could rise awareness for the future status of this species. The effects of climate change might be compounded by the geomorphology of the Adriatic Sea which could act as a “cul-de-sac” preventing whiting from shifting its distribution to higher latitudes [76]. Although dissolved oxygen was recognised as an influent predictor of local whiting biomass density, its temporal changes seemed somehow less related to the local biomass density trends. However, if dissolved oxygen would decline in large areas (e.g. due to climate change, increased river runoff and increased primary production) below the estimated threshold of 5.5 ml L^−1^ it could potentially influence whiting large-scale distribution. However, the findings of this study must be seen in light of some limitations. Firstly, our analysis did not separate whiting in age nor in length classes but investigated the overall response of the population to environmental variables. Since this response is often stage or length-specific, additional studies should be performed to investigate the potential different effects of environmental variables on different length-classes of whiting. Secondly, climate change may have other effects on whiting than a mere shift in distribution. For instance, organisms can undergo modifications in their reproductive capacities, physiology, phenology, growth, and mortality, leading to changes both at individual and population levels [77]. Whiting remains a not well studied species in the Adriatic Sea. More research is needed to deepen the knowledge of this fish, such as inter- and intra-specific interactions, fishing effects and aspects of its reproduction (e.g., influence of environmental variables on spawning and egg/larvae survival and transport), but also some aspects of its physiology (e.g., tolerance for low oxygen level and for increasing temperatures, and possible coping mechanisms). However, the present study contributes to expanding the actual knowledge on the ecology of whiting in the Adriatic Sea, providing information which might also be used as a baseline for future studies.

## Acknowledgments

We would like to warmly thank all the MEDITS team who have contributed to the data collection and organization. We thank Max Lindmark for the critical discussions on the modelling framework.

