## Supporting Information for "Spatio-temporal patterns of whiting (*Merlangius merlangus*) in the Adriatic Sea under environmental forcing"

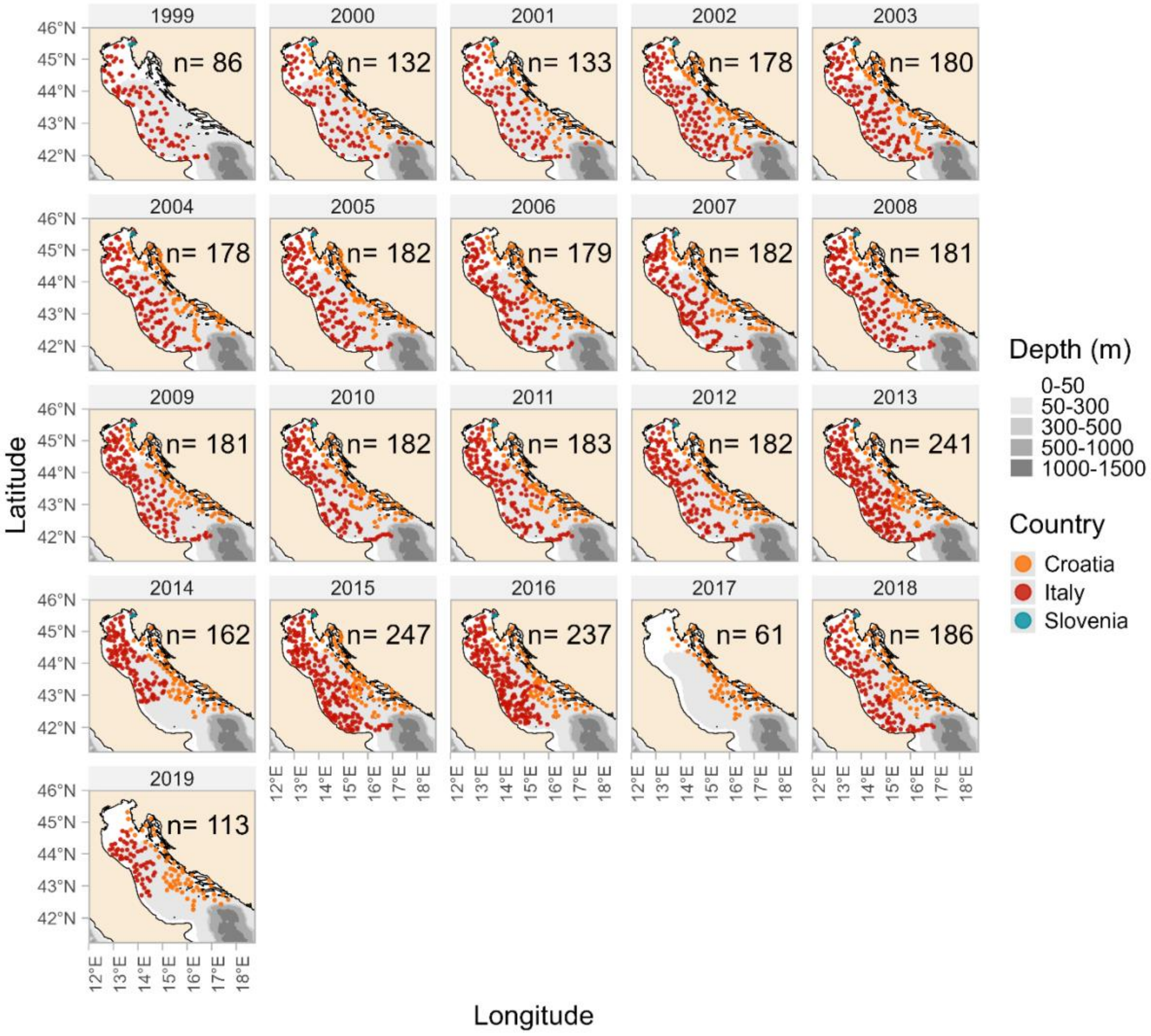


**S1 Fig. Trawl hauls locations per country.** The number of hauls conducted per year is reported on the top right of each panel. Missing hauls (e.g., in 2014, 2017, 2019) were conducted in autumn-winter period, thus removed from the analyses to obtain a more homogeneous dataset. In 1999 Croatia did not take part in the data collection.

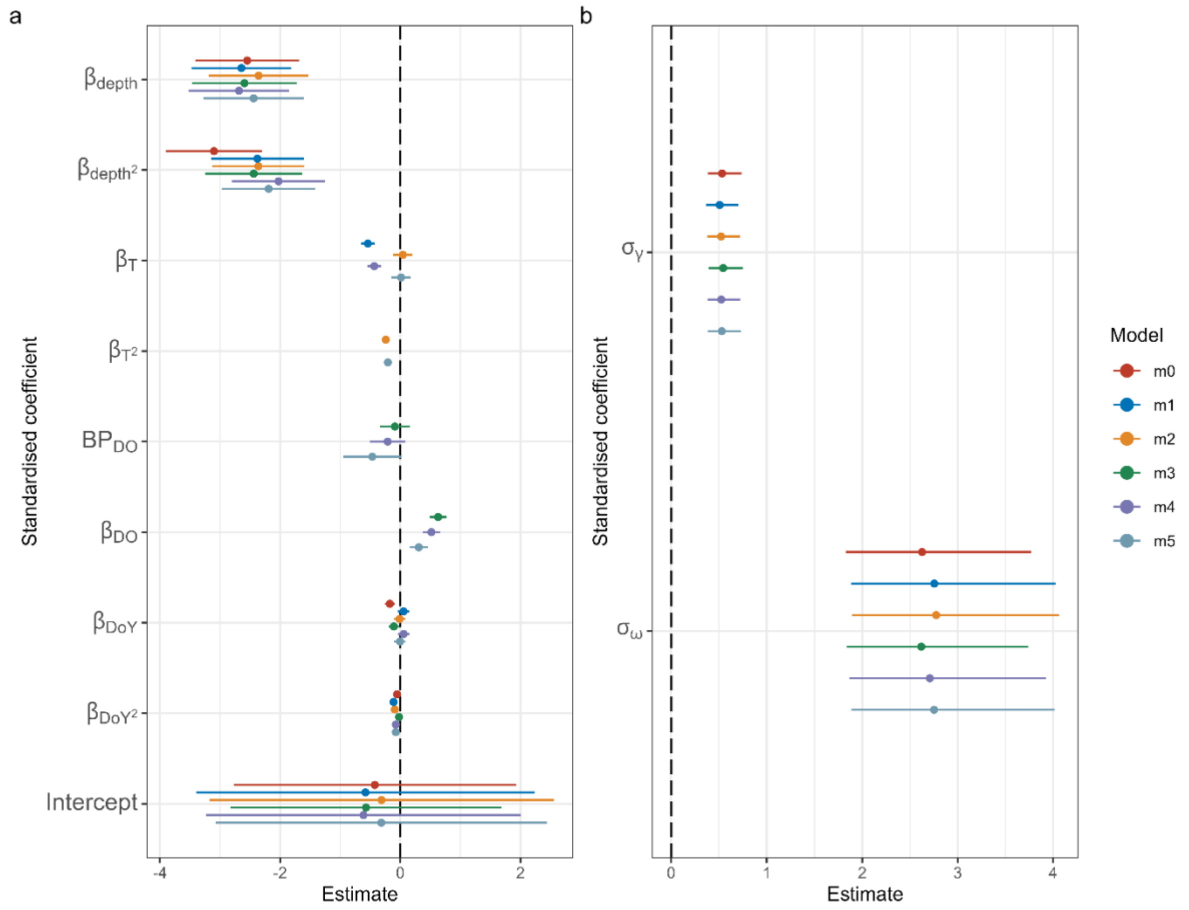


**S2 Fig. Standardized coefficients for the candidate models.** The point indicates the mean while the lines represent the 95 % confidence intervals for the fixed effects (a) and the random effects (b). $\beta_{DO}$ represents the coefficient of the slope related to DO while $BP_{DO}$ represents the estimated breakpoint. $\sigma_{\gamma}$ represent the standard deviation of the random year intercept and $\sigma_{\omega}$the standard deviation of the spatial random field.

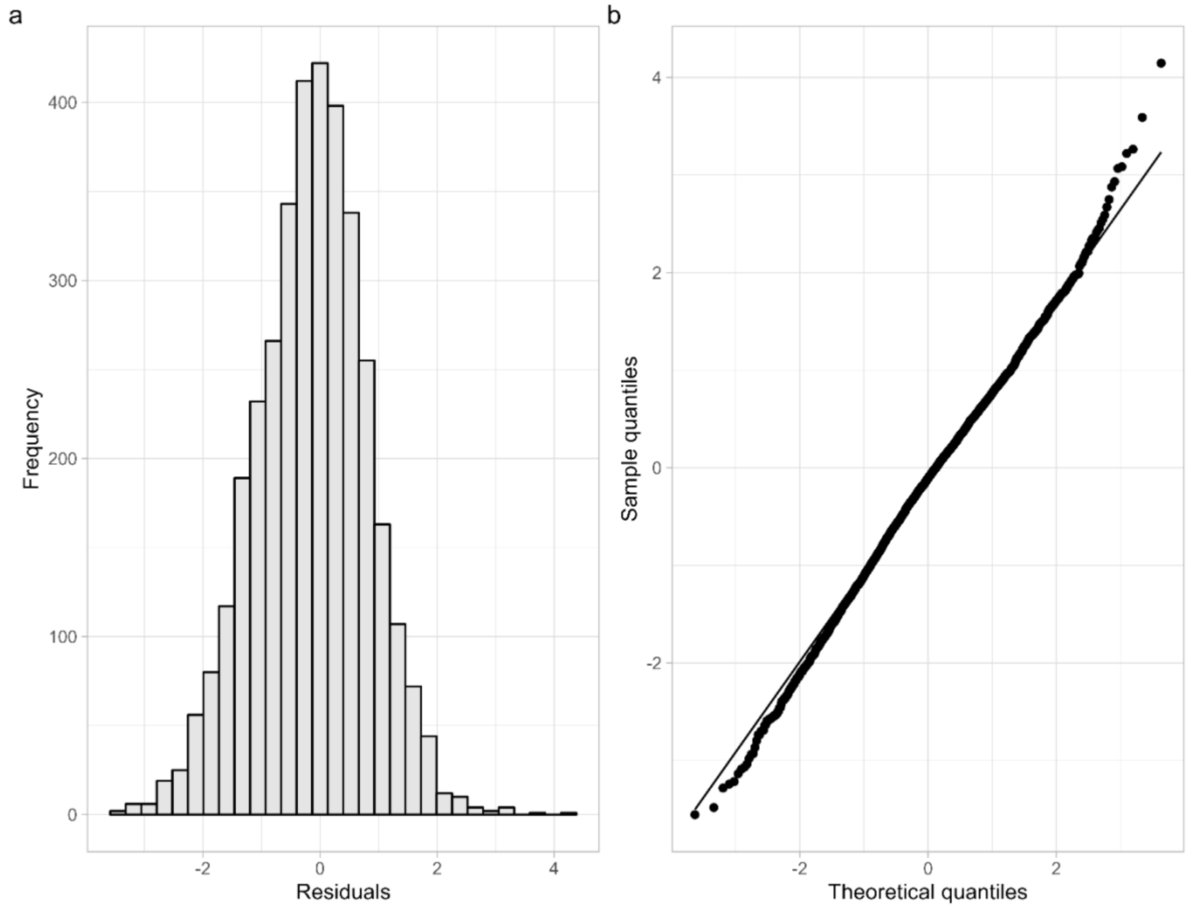


**S3 Fig. Randomized quantile residuals of the best fitting model.** Histogram of randomized quantile residuals (a) and residual quantile-quantile plot (b).

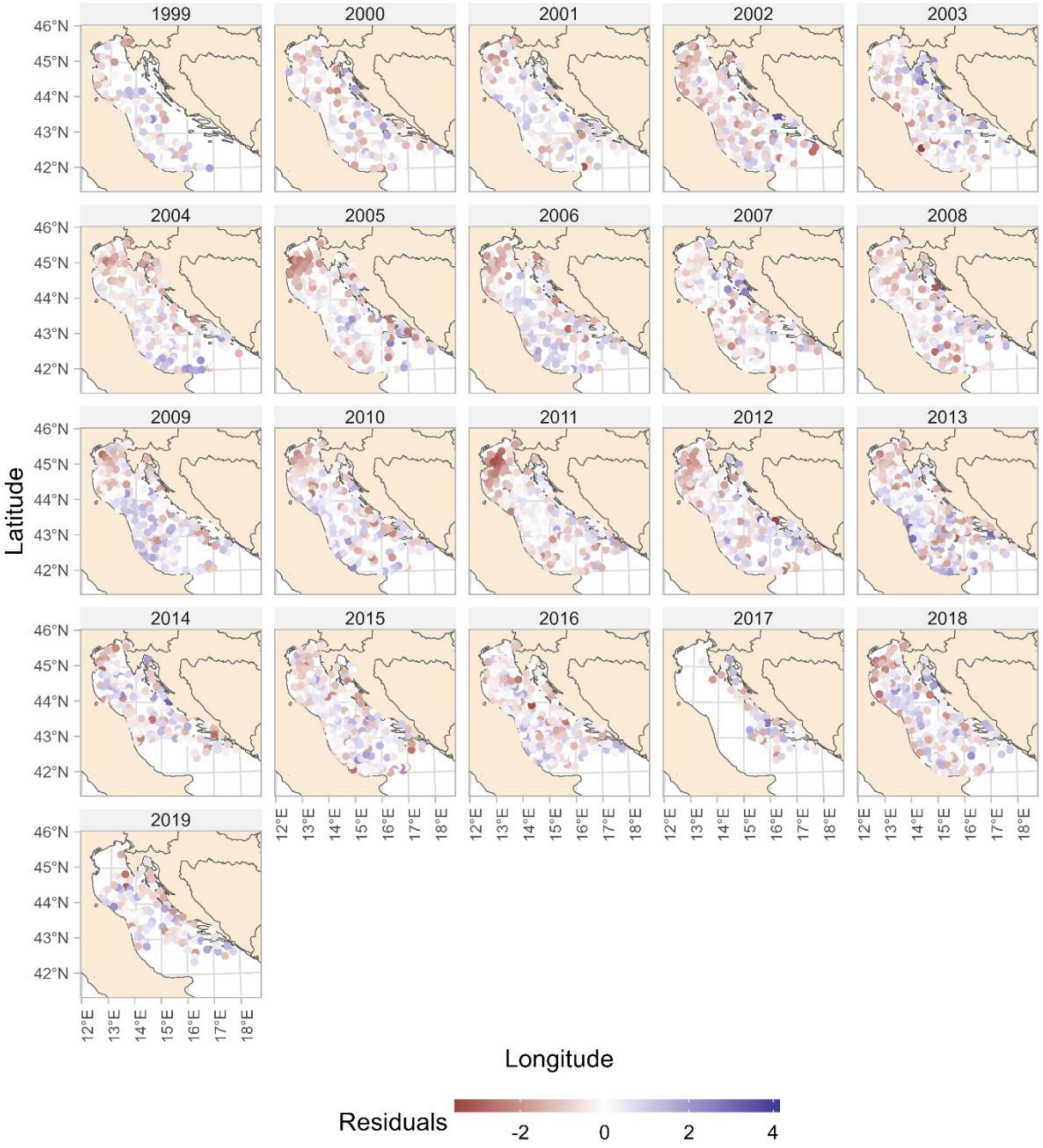


**S4 Fig. Quantile residuals plotted in space.**

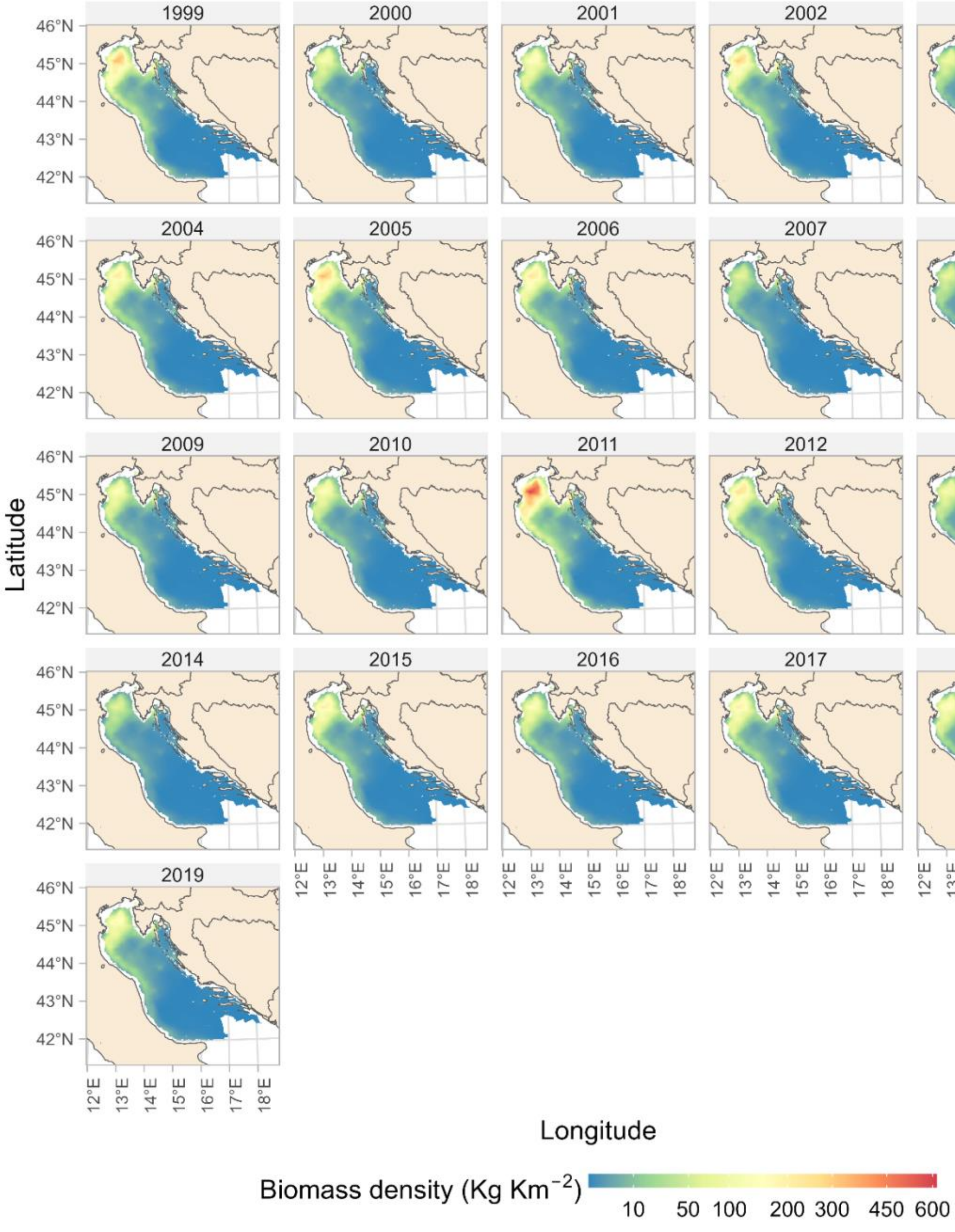


**S5 Fig. Predicted spatio-temporal changes of whiting biomass density (Kg Km$^{-2}$) by year.** All predictions were made over a 2 km$^2$ grid.
